# Linking visual and spatial exploration dynamics during free navigation in a large–scale virtual city

**DOI:** 10.64898/2026.04.06.714750

**Authors:** Vincent Schmidt, Debora Nolte, Jasmin L. Walter, Tracy Sánchez Pacheco, Peter König

## Abstract

Balancing exploration and exploitation is a fundamental challenge for adaptive behavior, yet it remains unclear whether visual sampling and spatial locomotion reflect a single cross–domain trait or operate independently. We addressed this question by recording head–mounted eye–tracking and full-body motion tracking while 26 participants freely navigated “Westbrook”, a large–scale virtual city for a total of 150 min across five sessions. From the movement trajectories we derived three spatial descriptors: median walking speed, occupancy entropy, and the proportion of explorative route choices. From the gaze data, we computed 38 robust visual descriptors encompassing fixation dynamics, pupil size, saccadic amplitude, gaze–head alignment, and transition entropy. Principal–component analysis reduced the visual descriptors to three components that captured 58 % of variance, with the first component (PC1) reflecting “gaze dynamism” (frequent shifts, larger saccades, higher transition entropy). Canonical correlation analysis revealed a strong coupling between spatial and visual behaviours: the first pair of canonical variates correlated at *r* = 0.68 (cross–validated *r* = 0.45), driven primarily by the association of high walking speed and occupancy entropy with elevated gaze dynamism. In contrast, the proportion of explorative route choices contributed little to this coupling. These findings demonstrate that individual differences in low–level locomotor speed and spatial coverage co–vary with an exploratory visual style, supporting the existence of a domain–general “exploration” factor that shapes both how people move through, and attend to, complex environments.

## Introduction

Across different domains of behavior, individuals must constantly balance the need to gather new information (exploration) with the need to utilize existing knowledge (exploitation) (Hills et al., 2015). In the visual domain, “visual explorers” are characterized by broad attentional sampling strategies and frequent gaze shifts, whereas “visual exploiters” concentrate their gaze on a limited subset of locations (Zangrossi et al., 2021). Similarly, in the spatial domain, individuals differ in how they traverse an environment: “spatial explorers” prioritize broad coverage and novel routes, while “spatial exploiters” (or returners) optimize known paths and revisit familiar locations (Pappalardo et al., 2015). Despite these striking conceptual parallels, a central question remains: do these behaviors reflect a unified, cross-domain trait? In other words, does an individual who broadly samples visual information also tend to explore physical space more extensively, or do visual and spatial exploration reflect independent dimensions of behavior (Kit et al., 2014)?

Answering this question has historically been difficult because these domains are typically studied in isolation. Consequently, classic paradigms (Olton & Samuelson, 1976; Tolman, 1948; Treisman & Gelade, 1980; Wolfe et al., 1989) are unfit to decouple the natural link between locomotion, head movements, and gaze (Foulsham et al., 2011; Tatler et al., 2011). To bridge this gap, experimental settings are required where visual and spatial behavior can be measured simultaneously under naturalistic conditions. For example, the use of immersive virtual-reality (VR) setups allows participants to move freely, engage in extended exploration, and interact with complex three-dimensional (3D) environments. As opposed to most real-world setups, it allows for precise tracking of all visible objects, including the participant’s head or body position, their gaze direction, and the name of the object hit by their gaze. Importantly, novel VR setups allow for eye and body movements to be recorded simultaneously under reliable, naturalistic conditions (Enders et al., 2021; Harris et al., 2022; Matthis et al., 2018).

Recent studies demonstrate that visual search traits are largely stable when comparing immersive and classic setups (Botch et al., 2023). However, they do not address how such individual differences relate to spatial exploration strategies during extended navigation, nor do they examine how visual and spatial behaviors jointly organize over time. This leaves a significant gap in our understanding of active perception: does the drive to sample novel information manifest as a domain-general trait (driving both gaze dynamics and locomotor patterns) or are these distinct control systems? For instance, it is plausible that high-level route planning operates under different metabolic and cognitive constraints than low-level saccadic targeting, potentially leading to dissociations between how an individual explores with their eyes as opposed to how they explore with their body. Consequently, it remains unclear whether individuals who broadly sample visual information also tend to explore space more extensively, or whether visual and spatial exploration reflect independent dimensions of behavior (Kit et al., 2014).

In the present study, we investigate this question by analysing whether distinct axes of visual exploration behavior covary with distinct features of spatial exploration. We test two complementary hypotheses using data from head–mounted eye– and location-tracking while participants freely explored a large-scale VR city (recorded in the context of Schmidt et al., 2023; analyzed in relation to spatial task performance in Walter et al., 2025). First, we hypothesized that participants who traverse the area more uniformly and faster (high spatial exploration) will also show richer visual dynamics, such as frequent gaze shifts and larger saccadic amplitudes. Second, we hypothesized that a small number of latent visual factors would capture the bulk of variance across high-dimensional gaze data, and that these factors would align with specific spatial strategies in a canonical correlation analysis (CCA).

## Results

We analyzed the eye-and body-tracking data of 26 participants who explored the immersive VR city “Westbrook” for a total of 150 min, divided across five 30–minute sessions. In the context of this free exploration, we measured how visually and spatially explorative or exploitative participants sampled their environment. From these recordings, we derived three spatial descriptors and summary statistics based on 11 visual features summarizing each individual’s explorativeness.

### Spatial exploration dynamics

The spatial descriptors summarize how spatially explorative each individual was. The first spatial descriptor aims to summarize how fast individuals walked and how much distance they covered as opposed to others. Given that each participant had the same time to explore, we calculated each individual’s median movement speed. The second spatial descriptor aims to summarize how unpredictably individuals moved through the environment. Occupancy entropy expresses the unpredictability of the individual’s transitions between areas within the environment. Higher entropy scores reflect less predictable movement patterns. The third and final spatial descriptor aims to summarize the explorativeness of individuals’ spatial decision-making. The proportion of explorative choices expresses how frequently individuals choose the less-frequented route when revisiting intersections. To provide examples, Figure 1 visualizes the three spatial features for a representative participant. The blue arrows show the individual’s updated locations for a 2-minute interval. It reveals a median movement speed of 9.85 kilometres per hour (km/h). The zoom-in on the top right shows the individual’s movement pattern, resulting in an occupancy entropy of 11.1, indicating a relatively uniform coverage of the area. At intersections, this participant chose the less–frequented branch 56% of the time, reflecting an explorative tendency. In combination, these features reflect spatial exploration dynamics.

**Figure 1.**
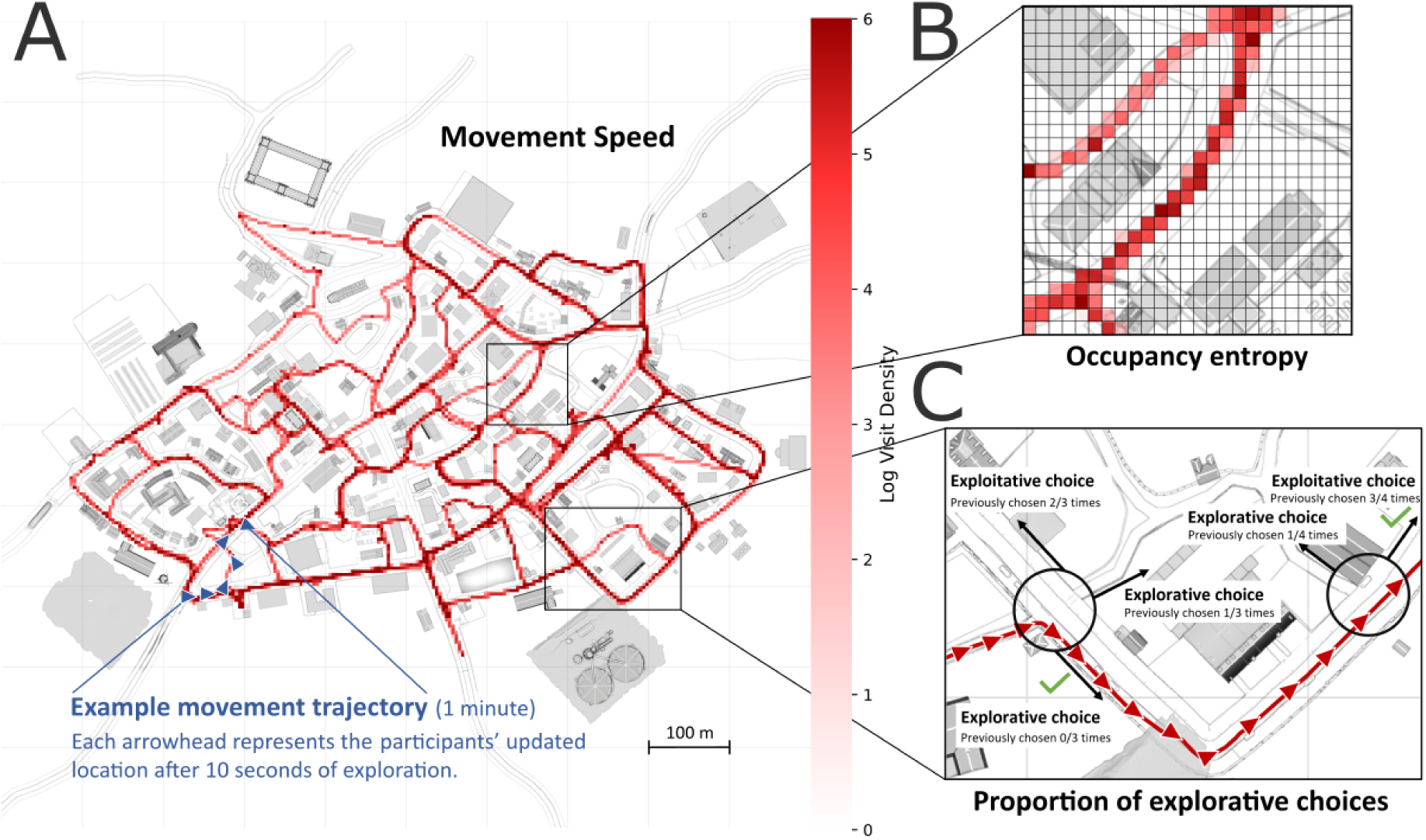
Spatial features. **(A)** The plot shows the log visit density of a single participant exploring the VR environment Westbrook (divided into 4×4m squares). The blue arrowheads exemplify the participant’s trajectory (2-minute segment) and heading direction, used to calculate movement speed. **(B)** To calculate occupancy entropy, we divided the map of the environment into 4×4m squares. The different shades of red overlaying the 4×4m squares show the log-transformed visit counts. **(C)** The proportion of explorative choices indicates how often the participant, when visiting an intersection (black circles), chose a path that they had chosen less frequently during previous visits than the other available options (explorative; i.e., participant’s choice at the second intersection) as opposed to the exploitative option (i.e., participant’s choice at the first intersection) that was visited more frequently, ignoring ties. In this example, the red line marks the participant’s movement trajectory and the red arrowheads indicate heading direction. The black arrows indicate possible route choices.

### Visual exploration dynamics

The visual features summarize how visually explorative each individual was. First, we accumulated a set of visual features based on the suggestions of previous research (Zangrossi et al., 2021) and expanded it to include a total of eleven features, each reflecting a different aspect of visual behavior. The first visual feature is the fixation duration, measured in seconds. The second visual feature is the number of fixations during 150 minutes of exploration. Third, the individual’s pupil diameter can indicate attention or arousal. It is measured separately for the left and right eye in millimetres. Fourth, gaze transition entropy is a value that expresses the unpredictability of transitions between different types of objects in the environment, with higher values indicating a more uniform distribution of transitions. Figure 2A provides visual examples for each of the visual features described above. Fifth, the distance from the observer to the fixated object in metres provides information on the visual depth of the individual’s gaze. Similarly, the distance between the targets of two consecutive fixations in metres reflects the visual depth of gaze transitions. Seventh, saccadic amplitude reflects the angular visual shift between consecutive fixations. Figure 2B provides a visual example of three aforementioned features. Eighth, the angle between gaze and head direction expresses angle between the direction of the gaze and the forward-direction of the head while fixating (see Figure 2C). Similarly, the angle between head-centered gaze directions of consecutive fixations expresses how individuals move their gaze, separately from head movements (see Figure 2D). Tenth, gaze shifts express how frequently individuals rotated their gaze in steps of 5° into either direction (see Figure 2E). By calculating shifts separately for rotations along the X- and Y-axes, as well as using head-and world-centered coordinates, we can investigate potentially different effects. Finally, the same separation applies to gaze flips (see Figure 2E), which are similar to shift but measure only rotational direction changes, exceeding the previously mentioned 5° threshold into the opposite direction as the reference. This set of visual features covers many aspects of explorative and exploitative visual behavior while the test subjects navigated the virtual environment.

**Figure 2.**
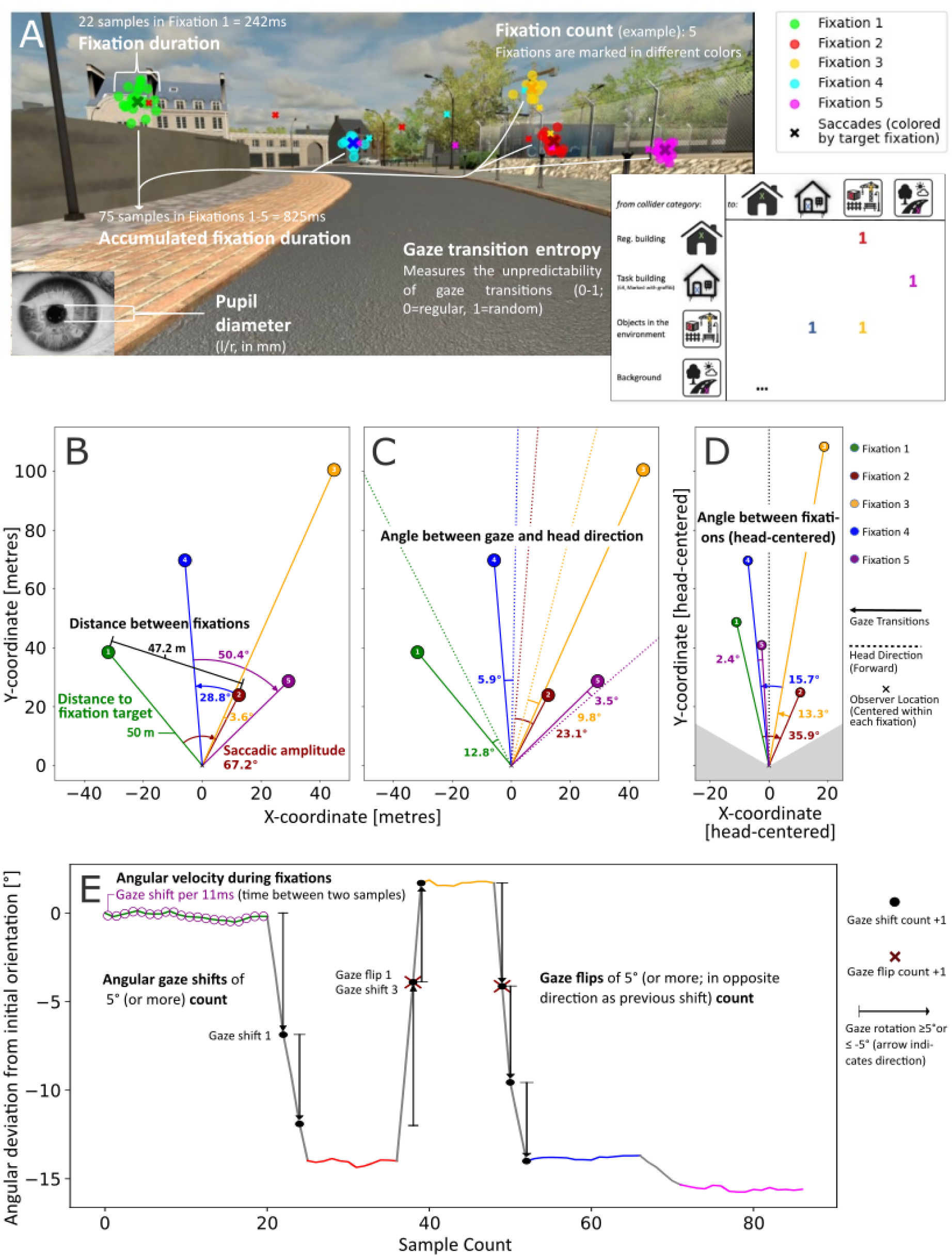
Visual Exploration Dynamics. (**A**) Example image of VR scene (from the perspective of the participant). The colored symbols show the gaze sequence of the example participant, divided into fixations (dots) and saccades (crosses) (see figure legend for sequence). The center of each fixation is marked with a large “X” in a darker tone of the same color. Importantly, the plot illustrates how (accumulated) fixation duration, fixation count, and pupil diameter were calculated. The inserted plot on the bottom right shows how transition entropy was calculated (unpredictability of transitioning between collider categories). (**B**) Bird’s-eye view of the visual scene described in panel A illustrating three measures assessing visual exploration dynamics: distance to the fixation (green), the distance between successive fixated objects (black), and saccadic amplitude (angle between the centers of successive fixations, red). (**C**) Shows the angle between gaze (solid lines) and head (dotted lines) direction during each of the example fixations. (**D**) Illustrates the head-centered angle between fixations, meaning the direction of the eyes normalized to the forward direction of the head, compared between subsequent fixations. (**E**) Illustrates how gaze shifts and gaze flips along the Y-axis were calculated (see Methods: Table 1). By applying the same procedure to the X-coordinate, and repeating it using head-centered coordinates, we can compare how frequently individuals rotated their gaze (world-centered) or their eyes (head-centered). The angular velocity during fixations expresses how much participants shifted their gaze between sequential individual samples (representing a time of 11 ms) within fixations. The result reveals how much individuals shifted their gaze while fixating.

After calculating all visual exploration features described above for each individual participant, we calculated measures of centrality, spread, and skewness for each continuous feature. These will be referred to as visual descriptors. Rather than relying on singular values (i.e., only centrality), which might be prone to errors, the set of resulting descriptors summarizes each individual’s distribution on each continuous feature. To avoid potential carry-over effects of outliers or measurement errors (Wilcox, 2012), we employed robust descriptive statistics. We calculated the median to measure centrality, the inter–quartile range (IQR) to measure spread, and Bowley’s coefficient of skewness (Sk). Figure 3 visualizes this procedure for the representative participant using their distance to the fixated object as an example. For the fixation duration, we additionally summed up all durations of individual fixations to receive the accumulated fixations duration (cf. Figure 2A). Finally, we separated the respective gaze data into head- and world-centered coordinates and further divided features related to gaze movements along their projected axes (horizontal, x-axis vs. vertical movements, y-axis). By calculating sums, median, IQR, and Bowley’s Sk, we derive a total of 38 visual exploration descriptors.

**Figure 3.**
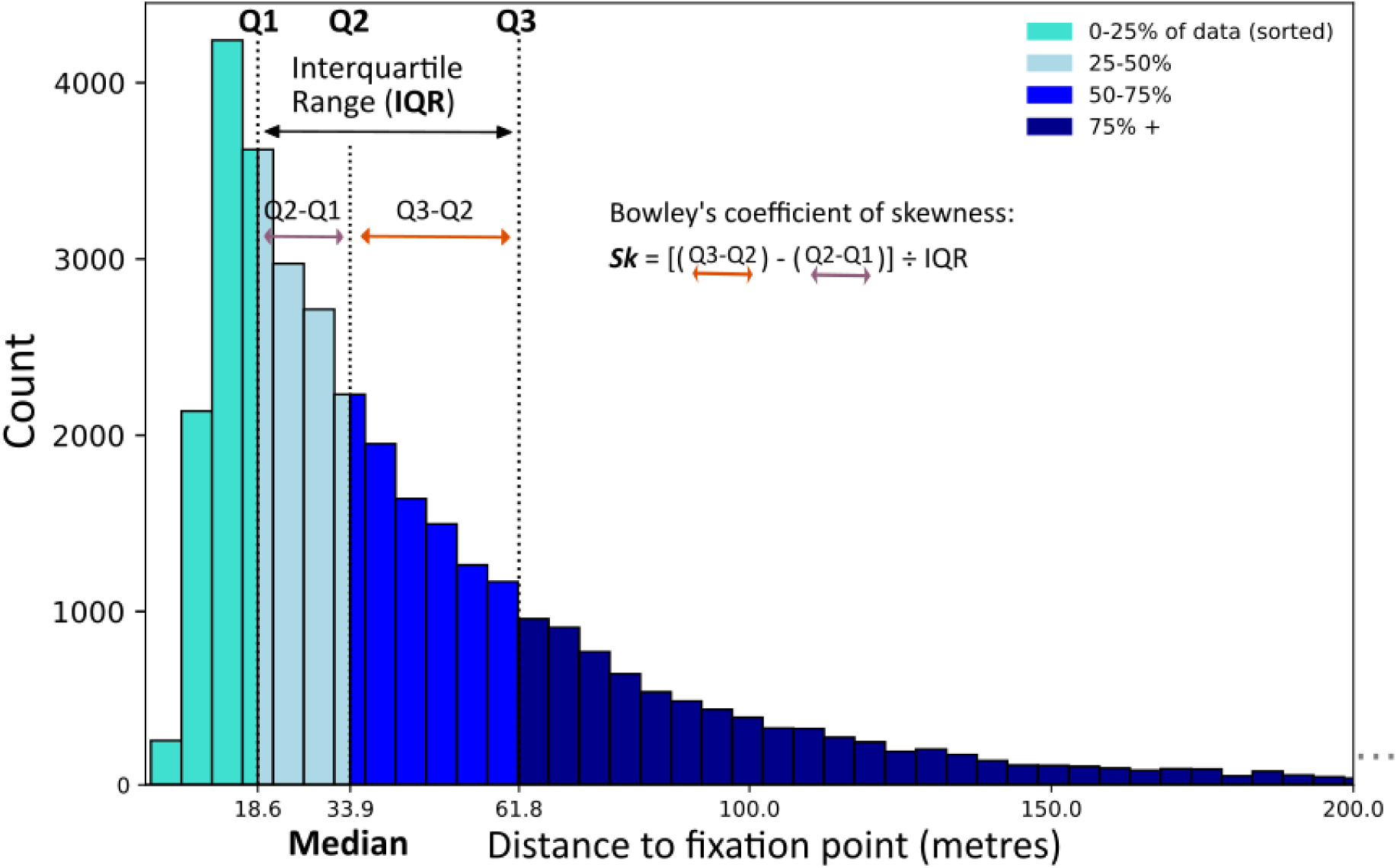
Example distribution (distance to fixation point). Visual explanation of IQR, quartiles, and Bowley’s coefficient of skewness (Sk) for a single participant. The plot shows the binned frequency counts for the distance between observer and observed object for the representative example participant (same as in Figure 1). The different shades of blue represent the different quantiles (see Figure legend). The arrows mark the IQR, which is the distance between the first and third quartile (black), the distance between the first and second quartile (magenta), and the distance between the second and the third quartile (orange). For each individual’s distribution on each continuous feature we calculated the median (Q2), IQR, and Bowley’s Sk.

### Comparing descriptors of spatial and visual exploration dynamics

First, we examined how the three spatial descriptors interrelated (Figure 4A). Movement speed correlated positively with occupancy entropy (*r* = .73, *df* = 24, *p* < .0001) and negatively with the proportion of explorative choices (*r* = –.67, *df* = 24, *p* < .001). Occupancy entropy and explorative choices were weakly anti–correlated (*r* = –.27, *df* = 24, *p* = .17). Thus, faster walkers tend to cover space more uniformly but make fewer intentional detours (novel route choices) at decision points.

**Figure 4.**
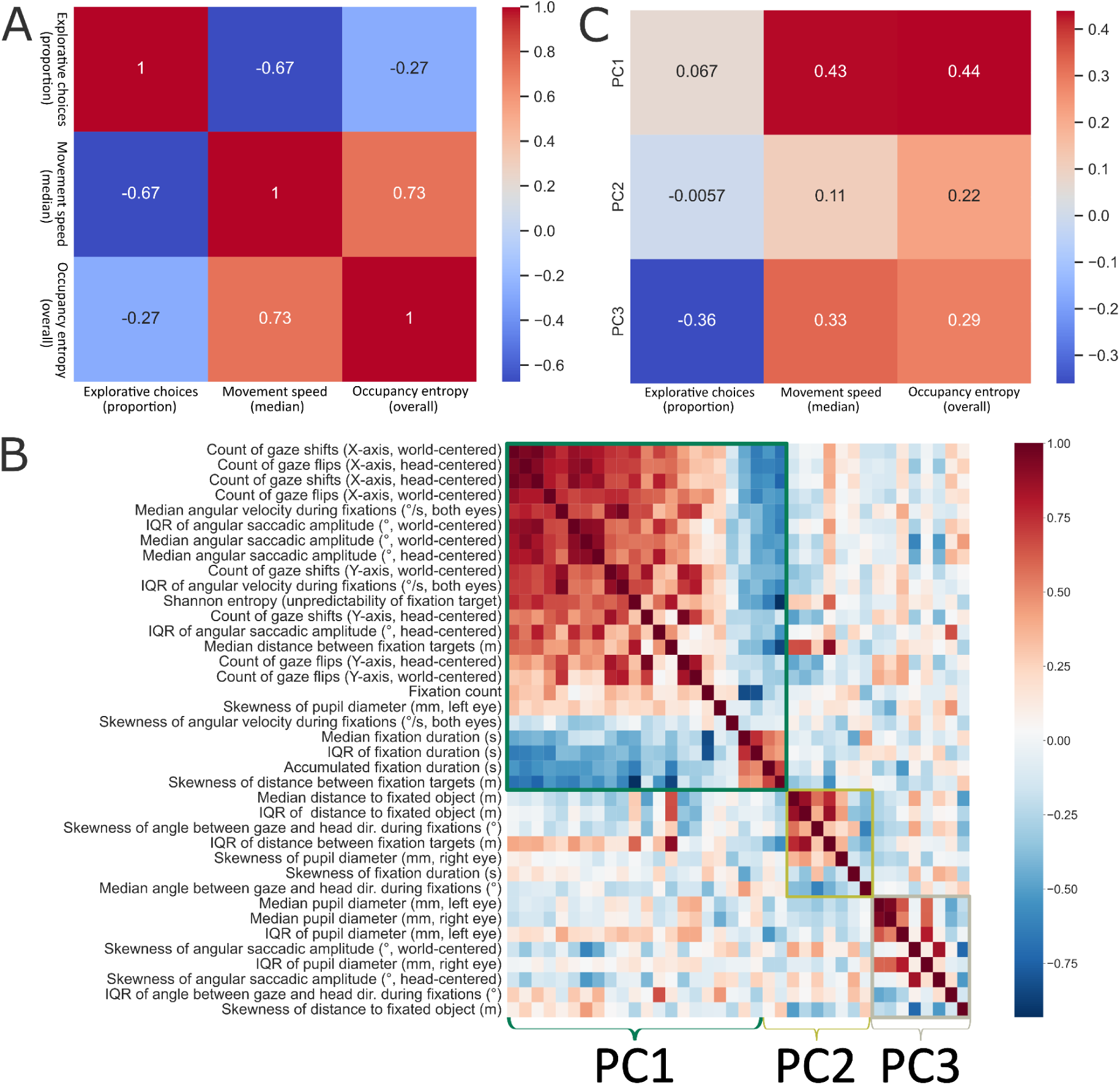
**(A)** Correlation Matrix of Spatial Descriptors. The heatmap and annotations show the Pearson correlation coefficients between the proportion of explorative choices, movement speed, and occupancy entropy. **(B)** Correlation Matrix of visual descriptors. The heatmap is showing Pearson correlations between all 38 descriptors addressing viewing dynamics. Descriptors are sorted by their loading on the respective PCs into a square matrix. The descriptors and their order are the same on the ordinate and abscissa. **(C)** Correlation Matrix between the three highest visual components (ordinate) and the three spatial descriptors (abscissa). Pearson correlation coefficients are annotated and reported using a heatmap.

To reduce the dimensionality of the 38 visual descriptors, we performed a Principal Component Analysis (PCA). The first three components together explain 58% of the total variance (Figure 4B). PC1 (33.1% variance) loads heavily on measures of gaze dynamism (frequency of shifts/flips, saccadic amplitude, angular velocity, transition entropy, fixation count, changes in eye-head alignment) and left–eye pupil–size skewness. It shows negative loadings on fixation–duration statistics. PC2 (15.3% variance) captures variability in fixation–to–object distance and inter–fixation distance (depth of visual search), as well as the angular deviation of the gaze from the head direction during fixations (head-centered gaze). PC3 (10.1% variance) is dominated by pupil–diameter centrality measures (left and right eye) and various measures of skewness (saccadic amplitude, angular velocity, fixation-to-object distance). Figure 4B displays the loading matrix and correlation heat–map among the original features. In essence, this analysis distills the complex array of eye-tracking metrics into three distinct behavioral profiles: how dynamically participants scan the scene (PC1), the depth at which they choose to focus (PC2), and their overall physiological arousal (PC3).

As a first step to investigate our core hypothesis, we determine the relationship between the first three principal components and the three spatial descriptors (Figure 4C). The resulting Pearson correlation coefficients (*df* = 24) indicate that PC1 (Gaze Dynamism) shows moderate positive correlations with occupancy entropy (*r* = .44, *p* < .05) and movement speed (*r* = .43, *p* < .05) and no significant correlation with the proportion of explorative choices (*r* = .07, *p* = .74). PC2 (visual depth) shows no significant correlations with movement speed (*r* = .11, *p* = .59), occupancy entropy (*r* = .22, *p* = .27), and the proportion of explorative choices (*r* = -.01, *p* = .98). PC3 (arousal) is not significantly correlated with the proportion of explorative choices (*r* = -.36, *p* = .07), neither with movement speed (*r* = .33, *p* = .1) or occupancy entropy (*r* = .29, *p* = .15). Gaze dynamism and visual depth increase together with occupancy entropy and movement speed and appear unaffected by explorative decision-making while physiological arousal appears to be additionally affected by it.

### Linking spatial and visual exploration dynamics

To test the core hypothesis of the study, the coupling between spatial and visual modes, we performed a canonical correlation analysis (CCA) between the visual PCs and the spatial descriptors. The CCA (Figure 5A) confirmed the relationships indicated by the correlation analysis (cf. Figure 4). The first pair of canonical variates captured a correlation of *r* = .68 (*df* = 24, *p* < .001) on the full dataset. To obtain a realistic estimate of generalizability, we applied leave–one–out cross–validation (LOO–CV), resulting in a cross–validated canonical correlation of *r* = .45 (*df* = 24, *p* < .05; Figure 5A). Redundancy analysis indicated that the first spatial canonical variate explains approximately 15.3% of the variance in the visual PC space, whereas the visual canonical variate explains approximately 17.7% of the variance in the spatial factor space. The cross-validated canonical correlation indicates a robust positive relationship between visual exploratory behavior and spatial exploratory behaviour across subjects.

**Figure 5.**
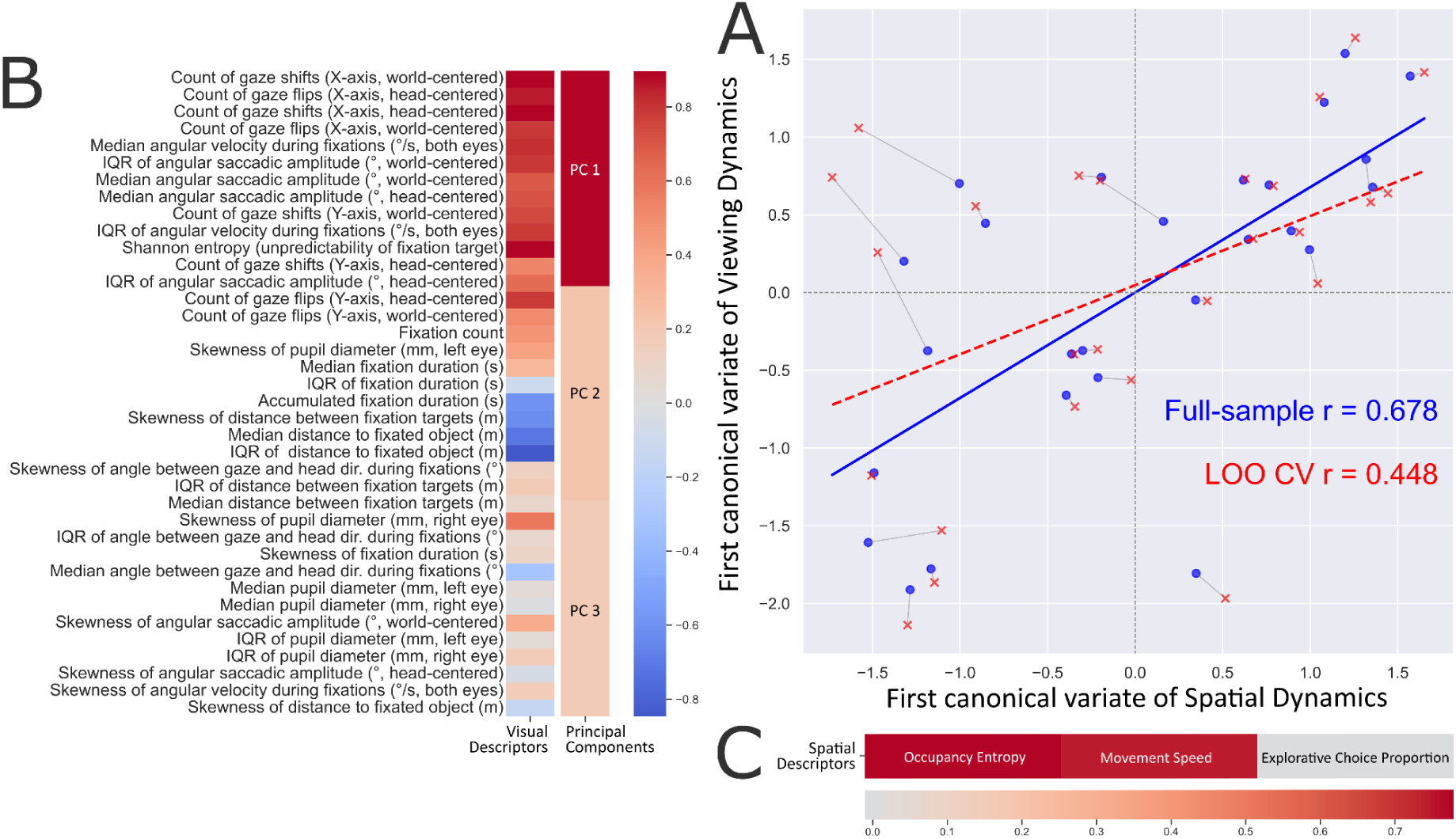
**(A)** Canonical Correlation between the first canonical variate pair. The scatterplot shows a canonical correlation of .68 between the first variates of visual and spatial dynamics for the full-sample (blue), and a canonical correlation of .45 after applying LOO-CV (red), as indicated by the colored markers and regression lines. Grey lines connecting data points indicate they were recorded within the same participant. **(B)** Heatmap of the projected loadings of each visual descriptor on the first visual canonical variate. The heatmap displays the loadings of the 38 visual descriptors on the first canonical variate of viewing dynamics. For convenience, the visual descriptors are listed in the same order as in Figure 4B. The second column shows the loadings of the three highest PCs on the first canonical variate that were created based on the visual descriptors. Colors indicate the magnitude of loadings. **(C)** Heatmap of the projected loadings of each visual descriptor on the first visual canonical variate. The heatmap along the abscissa shows the loadings of the spatial features on the first spatial canonical variate. Colors indicate the magnitude of loadings.

Next, we investigated how the individual visual features contributed to the first canonical variate of viewing dynamics (Figure 5B). First, we investigated the loadings of the first three PCs on the visual canonical variate. The Principal Components columns in Figure 5B shows that the first PC contributes the most to the visual canonical variate (loading = .95). The second (loading = .23) and third PCs (loading = .19) also contribute positively, though to a lesser degree. Then we projected these results back to the visual descriptors, considering their loadings on the three aforementioned PCs as well as their aforementioned loadings onto the canonical variate. The Visual Descriptors column reveals that, for example, the count of world-centered gaze shifts along the abscissa of the visual field and transition entropy (loadings = .9) have the highest loadings on the first visual canonical variate. The highest negative loading is the IQR of the distance from the observer to the fixated object (loading = -.85). Next to the descriptors included in PC1, especially the skewness of the pupil diameter of the right eye (loading = .58), the skewness of the angular saccadic amplitude (loading = .36), and the median angle between gaze and head direction during fixations (loading = -.32) contribute to the visual canonical variate. Overall, the first canonical variate of visual exploration dynamics is affected mostly by the descriptors included in PC1, as well as a few descriptors included in the remaining PCs.

Similarly, we investigated the loadings of the spatial descriptors on the spatial canonical variate (Figure 5C). Median movement speed (loading = .74) and occupancy entropy (loading = .77) contribute most to the canonical variate. Notably, the proportion of explorative route choices contributed minimally to this canonical relationship. Given that CCA maximizes the correlation between datasets, these results highlight that the strongest shared variance is between PC1 and the speed/entropy pair, underscoring that participants who move faster and cover space more uniformly also exhibit more dynamic gaze patterns.

## Discussion

### The present study provides the first systematic quantification of how visual and spatial exploration dynamics co–vary when participants freely navigate a large–scale virtual city

Concerning our first hypothesis, we found a robust coupling between locomotion and gaze: participants who traversed the area more rapidly and with higher occupancy entropy also displayed richer visual dynamics, characterized by frequent gaze shifts, larger saccadic amplitudes, and greater eye–head misalignment. This spatial–visual coupling persisted after dimensionality reduction and cross–validated canonical correlation. Providing an answer to our second hypothesis, a small set of three principal components captured the majority of variance across thirty–seven visual descriptors. The first component (PC1) aggregates measures of gaze dynamism with a skewness index of pupil size, suggesting that temporal variability in both eye–movement kinematics and autonomic arousal covaries across individuals. Thereby, PC1 clearly has the semantics of an exploratory PC. Third, the proportion of explorative route choices at intersections, which can be considered to be a behavioral marker of deliberative local decision–making, contributed little to the canonical relationship with visual dynamics. The correlations between spatial features additionally revealed the deliberative (local) exploratory choice does not lead by necessity to a larger occupancy entropy of the whole area. These results suggest that individuals showing explorative tendencies in low-level visual sampling also tend to be more spatially explorative, while individuals who use more exploitative strategies in the spatial or visual domain also tend to be more exploitative in the other.

While the current study resolves several limitations faced by previous research, we also identified several challenges. The current sample of experimental participants consists mostly of students and more diverse cohorts are required to test the generalizability of the spatial–visual coupling, especially across age groups. Nevertheless, while the magnitude of this relationship might be affected, it appears unlikely that it would disappear entirely. In contrast, the current findings might be used to investigate a potential link between spatial-visual (de-)coupling and clinical tests. Establishing a norm-image of spatial-visual sensory coupling might help to assist with identifying deviating cases.

Additionally, while head rotation was physical, forward locomotion was simulated via a joystick while participants remained seated. Incorporating treadmills or room–scale tracking would allow us to disentangle vestibular cues from visual–motor integration. Nevertheless, previous research showed that the swivel-chair and controller combination renders similar spatial results and spatial knowledge as a physical walking setup (Riecke et al., 2010) so we would not expect the actual results to change. Moreover, the addition of physiological recordings such as heart–rate variability could help to isolate the role of physiological arousal in this process more clearly. Regardless, due to the small overall influence of the pupil size descriptors, we would not estimate it to affect the results significantly. In sum, our findings highlight a coherent axis of individual differences that links how people move through a 3–D environment with how they allocate visual attention within it.

Our results extend a growing body of literature on active perception that emphasizes the bidirectional influence of locomotion and gaze. Our findings align with the results of Matthis et al., (2018) who showed that increased walking speed drives larger looking-ahead distances and saccades. By allowing full–body rotation and joystick–controlled forward movement during active exploration, we demonstrate that these effects survive in an ecologically realistic VR setting, supporting the view that the sensorimotor loop is preserved across contexts (Ruddle & Lessels, 2009). While recent work by Botch et al., (2023) demonstrated that visual search traits differ reliably between individuals and generalize from the lab to real-world scenes, their focus remained restricted to the visual domain. Their research did not address how such individual differences relate to spatial exploration strategies during extended navigation, nor did they examine how visual and spatial behaviors jointly organize over time. The strong loading of gaze–shift frequency on PC1 parallels findings from free–viewing of natural scenes, where shift–rate predicts information–seeking behavior (Kim, 2009; White et al., 2019). Furthermore, the canonical analysis reveals that faster locomotion aligns with a more exploratory visual style, consistent with the hypothesis that movement speed modulates visual sampling density (Gibson, 1979). Moreover, in spite of methodological differences with previous research, for instance how recordings were handled and how many unique features were included, the current study confirms that visual exploration dynamics are low-dimensional (Zangrossi et al., 2021). We show this effect persists in a realistic 3D setting, when using a diverse set of unique features and deriving robust descriptors to describe each participant’s individual distribution. Despite these striking conceptual parallels, a central question remains: do these behaviors reflect a unified, cross-domain trait? Our data provide empirical support for the idea that “visual explorers” (Zangrossi et al., 2021) and “spatial explorers” (Pappalardo et al., 2015) share a common underlying driver (Hills et al., 2015), potentially a general “curiosity” (Gottlieb & Oudeyer, 2018) factor, although this link appears stronger for low-level movement dynamics (e.g. speed/entropy) than for high-level navigational choices (e.g. explorative path choices).

The use of robust descriptive statistics (median, IQR, Bowley’s skewness) mitigates the impact of outliers that are common in VR eye–tracking (e.g., brief loss of pupil detection). The set of visual descriptors, according to our knowledge, is the most honest and complete set of different features expressing visual exploration dynamics used in a study on visual and spatial exploration. It considers factors that are often ignored, such as visual depth or isolating the eye from head movement. Moreover, the nested leave–one–out cross–validation employed for the CCA provides an unbiased estimate of the canonical correlation. The reduction from 0.68 to 0.46 after CV is comparable to reductions reported for high–dimensional behavioral datasets (Helmer et al., 2024), offering a realistic benchmark for future studies. These factors, as well as the unified spatial–visual framework presented here, offer a template for future investigations of sensorimotor coupling in immersive environments, including clinical populations with navigation deficits (e.g., Alzheimer’s disease: Coughlan et al., 2018) or studies of sensory augmentation (Schmidt et al., 2023).

## Methods

### Participants

Twenty–six healthy adults (age 18–35 years) were recruited from the city of Osnabrück and the surrounding area. Participants received either a monetary reward or university–internal participation credits. All participants were fluent in English or German. The study was approved by the Ethics Committee of the University of Osnabrück (protocol code: 4/71043.5, date of approval: 5 January 2021) and all participants gave written informed consent in accordance with the Declaration of Helsinki.

### Experimental procedure

After successful completion of a movement and a control tutorial in the form of a parcour, participants individually explored the Westbrook virtual environment five times (separate occasions with a maximum of 2 working days in between) for 30 minutes. In order to ensure participants had a reason to explore, several buildings (distributed in a balanced way across the environment) were marked with street art (for a more detailed explanation, see Schmidt et al., 2023). The city was designed with one Unity unit equating to one meter, allowing us to discuss distances in meters. Distances and coordinates were tracked using unity units, which equivalates to metres. For simplicity, we therefore refer to it as metres (m). The virtual city “Westbrook” (including 236 uniquely collidable buildings) contained realistic street layouts but no cardinal–direction cues (e.g., shadows or sun position) to avoid biasing orientation.

### Experimental hardware and setup

All sessions were conducted with the same experimental configuration. The virtual environment was rendered on an Alienware desktop (Nvidia RTX 2080 Ti, Intel i9, 32 GB RAM). Frame rates largely remained stable at 90 fps, never dropping below 60fps. Participants wore an HTC Vive Pro Eye (106° × 110° FOV) with built-in eye-tracking. Participants were seated on a swivel chair without a backrest; body rotations were captured by a chest-mounted body tracker and the VIVE tracking system, and were translated into virtual heading changes. Forward motion was simulated with a joystick on the Valve Index controllers; maximum virtual speed was limited to 18 km/h.

### Preprocessing

Gaze, head, and body tracking variables were synchronized during the experiment runtime. In addition, a Unity raycast was performed in viewing direction to obtain viewed object variables, whereas the information of the two hit colliders closest to the participant were saved. All synchronized data was saved with a varying sampling rate that averaged to 90 Hz. Since the virtual environment contained a large number small objects and partially or even fully invisible colliders (e.g. trees, lampposts, invisible functionality colliders, mesh wire fences; see fixations 3 & 5 in Figure 2A), as a first step in the data pre-processing, the closest collider hit was replaced with secondary collider hits if, the first collider was (partially) invisible, or belonged to the background category and the second collider was part of the building and object list. Next, eye movements were classified by an adapted version of the REMoDNaV algorithm, classifying eye movements with a dynamic velocity threshold (Dar et al., 2021; Nolte et al., 2025). The adaptation incorporated a correction of the participant’s translational movement during eye-event classification. Specifically, the eye tracking data was cleaned based on both eyes’ openness (>= 0.05), pupil diameter (!= -1), and the combinedGazeValidityBitmask (==3). In addition, to address the varying sampling rate and improve the event classification, we extended the REMoDNaV adaptation with a resampling step, in which we resampled all continuous data variables to 90 Hz (Walter et al., 2025). During the resampling, we also interpolated the cleaned eye tracking data, unless the continuous missing data segment exceeded 250 ms. Furthermore, initial events were only classified once they exceeded a minimum duration threshold (Dar et al., 2021; Keshava et al., 2023; Nolte et al., 2025). Otherwise, the pre-processing and event classification follow the filtering and processing steps proposed by Nolte et al., (2025), resulting in classified fixation and saccade events.

### Feature calculation

#### Spatial features

The proportion of explorative choices was quantified using the graph-based movement analysis pipeline described by Sánchez Pacheco et al., (2025). The occupancy map (Figure 1) was binarized to distinguish traversed from non-traversed locations, isolating walkable paths. To maintain topological continuity, minor morphological preprocessing was performed, such as filling small isolated gaps within street segments that could otherwise fragment the graph. The binary map was skeletonized, reducing all traversed paths to a one-pixel-wide structure while preserving geometric layout and connectivity. A street network graph was extracted from this skeleton, with nodes representing intersections or decision points and edges representing connecting street segments. To enhance spatial fidelity, node boundaries at decision points were manually adjusted to align with the actual spatial extent of intersections in the virtual environment, as determined by their virtual coordinates. Participant trajectories were then mapped onto the environment’s graph representation to classify navigational decision strategies. Each coordinate sample was assigned to the nearest graph element, either a node or an edge, resulting in a sequence of visited elements. At each node visit, the prior visit count for the selected outgoing path was compared with the visit counts for all available alternative paths at that decision point. Choices were classified as exploratory when participants selected a less-visited branch, and as conservative when they selected a more-visited branch. The resulting proportion of exploratory route choices served as a positive indicator of deliberative spatial exploration. Movement speed was calculated by dividing the 150 minutes of exploration into 10-second intervals. For each of these 10-second intervals, we calculated how far the participant moved and transformed the result into a velocity. From the resulting distribution of movement speeds across 10 seconds, we calculated the median for each participant. A faster median movement speed served as a positive indicator of spatial explorativeness. Occupancy entropy was calculated by transforming continuous positional coordinates into a spatial occupancy map (Figure 1) by discretizing the environment into 4 × 4 m cells and calculating visit frequencies for each of these. To quantify spatial variability of body movement, we computed a 2D occupancy-based entropy measure using the participant’s virtual body position coordinates. Positions were projected onto the horizontal plane and discretized into a fixed grid covering the full experimental environment (871.90 m × 578.60 m), divided into 218 × 145 bins corresponding to 4 m × 4 m spatial cells. For each participant, we calculated the frequency with which body positions fell into each grid cell and converted this distribution into probabilities. Spatial occupancy entropy was then computed using Shannon’s entropy. Higher entropy values reflect a more even and spatially distributed spatial exploration of the environment, whereas lower values indicate movement was concentrated in fewer locations.

#### Visual features

The visual features and the corresponding descriptors are inspired by previous literature (Zangrossi et al., 2021). Some features required further processing or calculations while other features were obtained directly. For instance, pupil diameter was directly provided by the eye tracker and was only filtered for validity as described in Preprocessing. Fixation duration, the distance to the fixated object, and the angular velocity during fixations were all calculated using Nolte et al.’s (2025) adaptation of the REMoDNaV algorithm (Dar et al., 2021). These continuous features were processed further into descriptors as outlined in Figure 3 and for fixation duration we additionally calculated the accumulated duration by summing up all individual durations within each participant. Fixation count was calculated by counting the resulting number of fixations classified by the algorithm. All features were standardized before performing the PCA and, again, before performing the CCA. Table 1 provides a detailed overview of each visual feature including how it was recorded, a technical explanation, the interpretation of the feature in the scope of the study, and the resulting descriptors that were calculated for the respective feature. In order to account for the fact that some features are less established than the five described above, we also describe those features below in detail.

**Table 1.** Technical descriptions of visual exploration features.

| Name | Measurement unit | Description (for each participant) | Resulting descriptors | Interpretation of the feature |
| --- | --- | --- | --- | --- |
| Pupil diameter | Millimetres (mm) | Pupil diameter of the left & right eye (separately); data was recorded by the eyetracker at a mean sampling rate of 90Hz. | Median, Interquartile range (IQR), Bowley's coefficient | Larger pupil sizes indicate increased stress levels or physiological arousal. |
| Fixation count | Count | The number of complete fixations detected (with start and end). Fixations were defined using the REMoDNaV algorithm (Dar et al., 2021; Nolte et al., 2025) | Count | More fixations are an indicator of explorative visual behavior. |
| Fixation duration | Seconds (s) | The duration of each fixation (one value per fixation); classified using the REMoDNaV algorithm (Dar et al., 2021; Nolte et al., 2025) | Accumulated, Median, IQR, and Bowley's coefficient | Shorter fixation durations are an indicator of explorative visual behavior. |
| Transitional entropy | Score | The unpredictability of the fixation sequence. Here, transitional entropy expresses the unpredictability of the participant's transitions between different collider categories: regular buildings, task buildings (marked with street art), objects (i.e., benches, poles, lamps etc), and background objects (i.e., pavement, trees, sky) (Sánchez Pacheco et al., 2026) | Overall score | A larger transitional entropy value is an indicator of explorative visual behavior. |
| Distance to fixated object | Unity units (metres / m) | The average distance between player head position and the hitpoint of the gaze on the fixated object for each fixation (one value per fixation), calculated using the REMoDNaV algorithm (Dar et al., 2021; Nolte et al., 2025) | Median, IQR, Bowley's coefficient | Fixating objects on a larger spatial distance is an indicator of explorative visual behavior. |
| Distance between fixation targets | Unity units (metres / m) | The distance between the centered hitpoint of one fixation to the centered hitpoint of the next fixation and fixated object for each fixation (one value per fixation). | Median, IQR, Bowley's coefficient | A larger distance between the targets of fixations might indicate explorative visual behavior. |
| Saccadic amplitude (head-centered) | Degrees (°) | The (head-centered) angle between sequential fixations. We calculated the angles between (head-centered) gaze vectors belonging to sequential fixations to estimate the angular size of the saccade. | Median, IQR, Bowley's coefficient | A larger saccadic amplitude is an indicator of explorative visual behavior. |
| Angle between gaze and head direction during fixations | Degrees (°) | The average angular deviation between eye and nose direction vector calculated between all timepoints within a fixation. | Median, IQR, Bowley's coefficient | A larger angular deviation between gaze and nose vector is an indicator of explorative visual behavior. |
| Angular velocity during fixations | Radians (°) per second | The combined angular velocity in radians/second during fixation calculated per sample using the REMoDNaV algorithm (max. 1000 radians/sec) (Dar et al., 2021; Nolte et al., 2025). Before calculating the descriptors, we applied mean-centering to each fixation. | Median, IQR, Bowley's coefficient | A larger angular velocity during fixations might be an indicator for explorative visual behavior. |
| Count of (head-centered) | Count | The count of how often the participant's world- and head-centered gaze direction was rotated by 5° | Count/Nr. | More world- or head-centered gaze |
| gaze shifts (separately for X- & Y- axes) |  | (independently of the direction of the previous rotation) or more. We calculated angles between all valid samples using the head-centered gaze direction. These angles were then summed up until the threshold of 5° or -5° was reached. Finally, we counted how often the threshold was exceeded into either direction. |  | shifts along the abscissa or ordinate of the visual field are an indicator of explorative visual behavior. |
| Count of (head-centered) gaze flips (separately for X- & Y- axes) | Count | The count of how often the participant's world- and head-centered gaze direction rotated by 5° or more into the opposite direction as the previous rotation. We calculated angles between each valid sample. These angles were then summed up until the threshold of 5° or -5° was reached. For flips, we counted how often the sign of the resulting angles changes from positive to negative or the other way around. | Count/Nr. | More world- or head-centered gaze flips along the abscissa or ordinate of the visual field are an indicator of explorative visual behavior. |

First, to measure how predictable participants’ gaze shifts were between environmental categories by building a first-order Markov transition matrix and calculating overall transition entropy, weighted by the stationary distribution (Sánchez Pacheco et al., 2026). To account for potential bias due to sparse transitions, we additionally computed a Chao–Shen bias-corrected estimate of transition entropy which rendered the same results for our dataset. Higher normalized entropy values indicate more heterogeneous, less predictable gaze transitions and broader visual exploration, while lower values indicate more repetitive transition structure. Second, to characterize the spatial distance between consecutive fixations, we first computed the three-dimensional centroid of each fixation based on the hit point coordinates on the object surface. For each participant, session, and recording segment, all hit point samples belonging to the same fixation were aggregated, and the centroid was calculated as the mean x, y, and z coordinate across samples within that fixation. These centroids therefore represent the spatially averaged fixation location in 3D space. Subsequently, we computed the Euclidean distance between each fixation centroid and the centroid of the immediately following fixation within the same participant, session, and segment. This yielded a measure of spatial displacement between successive fixations, reflecting how far gaze shifted from one fixation to the next with higher values indicating broader visual exploration. Third, to quantify directional changes between consecutive fixations, we first computed fixation-level centroids of the 3D world- and head-centered gaze direction vectors. For each participant, session, and recording segment, all gaze direction samples belonging to a given fixation were aggregated, and the centroid was calculated as the mean vector components across samples within that fixation. These centroids represent the average gaze direction during each fixation in world- or head-centered coordinates. Subsequently, we calculated the angular difference between consecutive fixation centroids within the same participant, session, and segment. The angle between successive mean gaze vectors was computed, providing a measure of the magnitude of directional reorientation from one fixation to the next with larger angular reorientation indicating broader visual exploration. Fourth, to quantify fixation-level eye orientation relative to the head, we computed centroids of the local 3D eye direction vectors for each fixation. For every participant, session, and recording segment, all local gaze direction samples belonging to a fixation were aggregated, and the centroid was calculated as the mean vector components (x, y, z) across samples within that fixation. These centroids represent the average eye direction in head-centered (local) coordinates during each fixation. Subsequently, we calculated the angular deviation between each fixation’s mean local eye direction vector and a forward-facing reference vector (nose vector: [0, 0, 1]). The resulting angle reflects the extent to which the eyes deviated from the head’s forward axis during each fixation, providing a measure of eye-in-head eccentricity, which might be an indicator of visual explorativeness. Finally, gaze shifts were identified using an angular thresholding approach. A gaze shift was counted whenever the world-centred gaze angle exceeded a ±5° threshold relative to the current reference orientation. When the angle subsequently crossed the threshold in the opposite direction, this was counted as a flip, indicating a reversal in gaze direction. The same procedure was applied to head-centred coordinates to isolate gaze rotations occurring relative to the head rather than in world space. The resulting counts of gaze shifts and flips provide an index of directional instability and reorientation frequency, with higher values reflecting visual exploration and lower values reflecting visual exploitation.

### Statistical analysis

Principal component analysis (PCA) was performed on the z–scored set of 38 robust visual descriptors. Canonical correlation analysis (CCA) was conducted on the z-scored results of the PCA and the three spatial features, with a nested leave–one–out cross–validation (LOO-CV) to obtain unbiased estimates (see Data & Code Availability). Due to the specific design of the analysis, we focused our analysis entirely on the first pair of canonical components as this is by design the strongest correlation.

### Dataset and power considerations

The dataset constitutes the control group of Schmidt et al., (2023) where we compared the participants’ performance on a subsequent spatial assessment with an experimental condition. The preceding exploration sessions that are analysed in the current study were first published in Walter et al., (2025), investigating the relationship between individual differences and the relationship with spatial tasks. While an a priori power analysis was performed for the original between–group comparison, the sample size (*N*=26) provides sufficient degrees of freedom for the exploratory correlational techniques (PCA, CCA) employed here (Cohen, 1992).

## Data availability

All data, as well as analysis scripts were made publicly available. The raw data can be downloaded here: https://osf.io/qcn67/, and the preprocessing code is available here: https://github.com/JasminLWalter/VR-EyeTracking-GraphTheory/tree/master/GraphTheory_ ET_VR_Westbrueck/Pre-processing2023. The original scripts used for the calculation of transitional entropy are available here: https://github.com/tracysanchez/Human_Agents_Impact_Nav. The analysis code and the adjusted scripts employed in the statistical analysis of the current study are available here: https://osf.io/uxv8k/.

## Acknowledgements

We thank Kaya Gärtner and Aura Kampf for their support in selecting and calculating the spatial variables, specifically the proportion of explorative decisions. We thank Philip Spaniol for his support with the creation of the virtual environment and the maps. Finally, we thank Nora Maleki, and Linus Tiemann for their technical support as research assistants in the development of the VR environment.

## Informed consent

Informed consent was obtained from all subjects involved in the study.

